# Translational activity is uncoupled from nucleic acid content in bacterial cells of the human gut microbiota

**DOI:** 10.1101/2020.10.05.327163

**Authors:** Mariia Taguer, B. Jesse Shapiro, Corinne F. Maurice

## Abstract

**Background:** Changes in bacterial diversity in the human gut microbiome, characterized primarily though DNA sequencing methods, have been associated with many different adverse health conditions. However, these changes do not always reflect changes in bacterial activity, and thus how the gut microbiome is implicated in disease is still not often understood. New methods that link together bacterial function to bacterial identity are needed to further explore the role of the gut microbiome in health and disease. We optimized bioorthogonal non-canonical amino acid tagging (BONCAT) for the gut microbiota and combined it with fluorescently activated cell sorting and sequencing (FACS-Seq) to identify the translationally active members of the community. We then used this novel technique to compare and contrast to other methods of bulk community measurements of activity and viability: physiological staining of relative nucleic acid content and membrane damage. Relative nucleic acid content has previously been linked to metabolic activity, yet remains currently undefined for the human gut microbiota.

**Results:** Ten healthy, unrelated individuals were sampled to determine the proportion and diversity of distinct physiological fractions of their gut microbiota. The translationally active bacteria represent about half of the gut microbiota, and are not distinct from the whole community. The high nucleic acid content (HNA) bacteria also represent about half of the gut microbiota, but are distinct from the whole community and correlate with the damaged subset. Perturbing the community with xenobiotics previously shown to alter bacterial activity but not diversity resulted in stronger changes in the distinct physiological fractions than in the whole community.

**Conclusions:** BONCAT is a suitable method to probe the translationally active members of the human gut microbiota, and combined with FACS-Seq, allows for their identification. The high nucleic acid content bacteria are not necessarily the protein-producing bacteria in the community, and so further work is needed to understand the relationship between nucleic acid content and bacterial metabolism in the human gut. Taking into account physiologically distinct subsets of the gut microbiota may be more informative than relying on whole community profiling.

## Background

The human gut microbiota is comprised of trillions of microorganisms that together help maintain gut homeostasis and provide key ecosystem services to the human host such as nutrient degradation, pathogen exclusion, and immune system development[1, 2]. The diversity of the gut microbiome across different human populations and across various diseases has been well catalogued[3–8]. Still, it is becoming increasingly clear that bacterial diversity and activity are not always coupled, and thus which members are key for maintaining host health cannot be elucidated through diversity metrics alone[9–11]. New methods that link together bacterial function to bacterial identity are needed to further explore the role of the gut microbiome in health and disease.

The active subset of the gut microbiota is more sensitive and responsive to perturbations than the diversity of the whole community alone[12, 13]. Broad range ‘omics techniques such as metatranscriptomics, metabolomics, and metaproteomics provide an overall depiction of the gut microbiota’s output, yet incomplete functional databases make it challenging to link together bacterial identity to function[14]. To characterize the active fraction of the gut microbiota, a broad and measurable indicator of activity needs to be identified that labels the active bacteria with minimal bias. Single-cell techniques allow for increased resolution of the heterogeneity of activity in complex microbial systems, helping determine the actual contribution of specific bacterial members *in situ*. Heavy water incorporation, substrate uptake, and nucleic acid content have all been studied in the context of the gut microbiota[10, 12, 15–18]. Yet they have low throughput, are costly, or, for the case of nucleic acid content, the relevance to activity is unclear. Single-cell techniques to study bacterial physiology and activity have been well-detailed by Hatzenpichler *et al* (2020)[19].

The use of relative nucleic acid content as a marker of bacterial activity was first introduced in aquatic systems, where the microbial community, when stained with a nucleic acid dye, clusters into two distinct populations based on their level of nucleic acid content[20]. The more fluorescent population consists of bacteria with higher nucleic acid content (HNA) than their low nucleic acid (LNA) counterparts. This phenomenon has since been widely studied, and is proposed to link together bacterial nucleic acid content to a gross level of bacterial metabolism, where the HNA bacteria are more metabolically active than the LNA. This has been demonstrated through higher leucine incorporation rates, ATP cell^-1^ concentrations, respiration rates, and proportions correlating with overall bacterial production [21–27], but these dynamics have been disputed as well [28].

As this bimodal distribution of HNA and LNA has already been identified in the gut[10], we set out to determine if HNA and LNA components of the microbiota differ in their metabolic activity. To do so, a broad yet clearly defined measurement of single-cell levels of activity was required to compare and contrast with relative nucleic acid content determination. In this paper, we focus on protein translation as an important aspect of metabolic activity. We optimize bioorthogonal non-canonical amino acid tagging (BONCAT), a recent application of click chemistry, to identify the translationally active bacteria in the gut microbiota[29]. BONCAT allows for the unbiased detection of proteins produced *in situ* under biologically relevant conditions, without the need for radioactivity, isotopes, antibodies, long incubations, or altering conditions. A methionine analogue, L-homopropargylglycine (HPG), is added to a short *in vitro* incubation of the gut microbiota, and is then “clicked” to an azide-modified fluorophore. HPG has been shown to be taken up by all bacteria under all physiological states tested, and due to the promiscuity of methionyl-tRNA synthetase, is incorporated into nascent proteins[29]. The alkyne-azide groups quickly undergo a cycloaddition to form a stable triazole conjugate at biologically relevant conditions. Azide and alkyne modifications are considered biologically inert: they do not interfere with biological processes and do not naturally exist in most biological systems, including bacteria[29–31].

Previously, BONCAT has been used in bacterial isolates, natural assemblages in aquatic systems[30–32], soil[33], and sputum from cystic fibrosis patients[34], and typically combined with fluorescent in-situ hybridization (FISH) with 16S rRNA probes to identify the protein-producing bacteria. To our knowledge, ours is the first study to apply BONCAT to the gut microbiota. As 16S rRNA-FISH offers limited taxonomic resolution and requires designing probes for specific taxa of interest *a priori*[35], we instead combine BONCAT with fluorescence-activated cell sorting (FACS) and subsequent 16S rRNA gene sequencing (FACS-Seq) as previously done elsewhere[33, 34, 36]. This allows us to identify the diversity of the BONCAT+ and BONCAT- communities, linking together bacterial identity to activity and increasing throughput. Assessing protein production through BONCAT yields similar results to assessing protein production through nano-SIMS^[29]^and MAR-FISH^[30]^, yet it is faster and less expensive.

Applying BONCAT to the gut microbiota provides some unique challenges that we address in this study. The incorporation of HPG into nascent proteins requires a short *in vitro* incubation, which is difficult for the gut microbiota, as there is no single media that is able to support the growth of all gut bacteria. Indeed, “culturomics” is an ongoing, developing field to broaden our ability to culture more of the fastidious members of the gut microbiota[37]. Furthermore, methionine is common in the gut, but inhibits HPG incorporation. HPG has an activation rate 500 times lower than methionine[38], and bacteria preferentially incorporate methionine over HPG ten-fold, so excess HPG is required to outcompete methionine[30]. Methionine is expected to be ubiquitous in the gut lumen, and as such, fecal bacteria are not immediately well-suited for BONCAT. The approach detailed in this study addresses and overcomes some of these limitations.

Lastly, to explore what the inactive subset of the gut microbiota may represent, we identified the damaged subset of the gut microbiota with propidium iodide (PI), a membrane exclusion dye. By contrasting the active subset to the damaged subset, we hope to begin to explore if the less active or inactive fraction represent dormant bacteria acting as a seedbank, or external transient bacteria that are unable to colonize the gut[39–43].

Here we show that the HNA and BONCAT+ bacteria are taxonomically distinct, suggesting that the HNA bacteria are not necessarily undergoing translation. We explore how the HNA bacteria contain more of the conserved bacteria across individuals, contributing to the “core” gut microbiome, while the BONCAT+ community is a subset of the whole community and potentially more responsive to changes in the environment.

## Methods

### Sample collection

Human studies were performed with approval of the McGill Ethics Research Board (REB #A04-M27-15B). Ten healthy, unrelated individuals who had not taken antibiotics in the past 6 months and had not been diagnosed with a gastro-intestinal condition provided fecal samples on site. Samples were immediately placed in the anaerobic chamber (Coy Laboratory Products, 5% H_2_, 20% CO_2_, 75% N_2_). Sample preparation and staining were performed in the anaerobic chamber; FACS was performed aerobically. Metadata questionnaires were completed post-sample donation, collecting information on dietary logs for the previous 48 hours, as well as recent travel history, antibiotic usage, and typical coffee, chocolate, tea, and dairy consumption.

### Gut microbiota sample preparation

A portion (1-4 g) of the fresh fecal sample was diluted 1/10 w/v in reduced PBS (rPBS) containing 1 ug.ml^-1^ resazurin sodium salt and 1 mg.ml^-1^ L-Cysteine (final concentrations). Samples were thoroughly vortexed and centrifuged at 700 x *g* for 1 minute to pellet large organic debris. Supernatant was washed 3 times with rPBS by centrifuging for 3 minutes at 6,000 x *g*, removing supernatant, and resuspending with fresh rPBS. Samples were then diluted in rPBS for staining and FACS. Staining was performed anaerobically in the dark with 1X SYBR Green I (Invitrogen) for 15 minutes, or PI (Sigma-Aldrich) at 0.04 mg.ml^-1^ for 10 minutes[44].

### BONCAT incubations, growth curves, and click reaction

Bacteria were diluted 1/10 in 50% (v/v) of the supernatant retained from the first 6,000 x *g* centrifugation, 2mM final concentration HPG, and remaining volume of rPBS. Bacteria were incubated at 37°C for 2 hours unless otherwise stated. When specified, glucose or drug additions were added at the start of the BONCAT incubations. Glucose was added at a final concentration of 0.2%, and nizatidine and digoxin added at 0.01 mg/ml. A no-HPG incubation is included as a control, and each sample is incubated in duplicate (triplicate for the xenobiotic experiments). To stop the incubation, bacteria were fixed with 80% ethanol to a final concentration of 50% (v/v). Samples were stored at 4°C until processed with the click reaction that same day.

For the click reaction, bacteria were pelleted and resuspended in the click reaction solution (Click-iT Cell buffer kit, ThermoFisher Scientific) containing 5µM Alexa-647 azide, and incubated in the dark at room temperature for 30 minutes. A no-Alexa control is included. Samples were then centrifuged at 8,000 x *g* for 5 minutes, supernatant removed, and washed with 80% ethanol, and then centrifuged again and resuspended in PBS and stained with SYBR Green I. Growth curves were performed anaerobically in the dark in 96 well plates, with triplicates for each condition, using the BioTek Epoch 2 microplate spectrophotometer at OD_600_ with shaking before each measurement.

### Fluorescence-activated cell sorting (FACS) and cell counts

Cell sorting was performed on the FACSAria III (BD Bioscience) equipped with a 488 nm laser and the appropriate detection filters. Positively stained cells were determined from debris and unstained cells using unstained controls. 180,000 events were sorted using a 70 μm nozzle for each population for each individual and frozen at -80 °C for later DNA extraction. Sheath fluid was collected at the end of every sorting day as a negative control to detect contaminant DNA. Data files were analyzed using FlowJo V7 software (FlowJo LLC). Cell count data was analyzed as previously reported[45].

### DNA extraction and 16S gene amplicon bioinformatics analysis

Samples were stored at -80°C until DNA extraction. Samples were extracted with the Qiagen AllPrep PowerFecal kits as per the manufacturer’s instructions. The V4-V5 hypervariable region was amplified with the 515F/926R primers[46]. Trimming, alignment of paired end reads, and quality filtering was performed by DADA2[47]. Taxonomic alignment was performed with a pre-trained Naives Bayes classifier using SILVA 132 database on 99% OTUs using QIIME2.

Taxonomic and low abundance filtering was performed in phyloseq (v1.3) in R (v3.6.1). Reads present in the sheath fluid but absent in the whole community samples (69 ASVs) were removed. As well, ASVs without phyla-level taxonomic assignment were removed. A prevalence threshold was set at a minimum of 6 reads in at least 2 samples. Finally, two *Pseudomonas* ASVs were identified as contaminants and removed.

Count data was rlog transformed using the DESeq2 package (v1.26) and weighted UniFrac distance matrix was calculated using rbiom (v1.0). Beta diversity was assessed on weighted UniFrac distances using pairwise PERMANOVA with 999 permutations to test for significance using adonis in the vegan package (v2.5). Differential abundance testing was performed using the statistical analysis package corncob (v0.1) which performs beta-binomial regression models to determine differentially abundant and dispersed relative abundances. Weighted UniFrac distances between physiological groups were compared using the Kruskal-Wallis test in the rstatix package (v0.5). Alpha diversity of the sorted and unsorted comparisons was performed on samples rarified to 10534 reads/sample without replacement using the Shannon diversity metric.

## Results

### Optimizing incubation conditions for HPG labelling of gut bacteria

The BONCAT method requires an incubation step as bacteria incorporate HPG. In the interest of maintaining gut bacterial activity to levels as similar to *in situ* as possible, optimization of the media, length of incubation, and concentration of HPG was necessary to ensure that bacterial growth and community structure was not altered. To best mimic the gut environment, a fecal slurry was created by homogenizing fresh stool in reduced PBS (rPBS – 1 mg mL^-1^ L-cysteine), and the supernatant of the fecal slurry was tested as the media at varying concentrations, along with different incubation times. An additional challenge to an *in vitro* BONCAT incubation is that methionine is preferentially incorporated over HPG[29, 30], and the gut is a nutritionally rich environment with high concentration of methionine **(Table S1)**. As such, HPG was added in excess of 10X what is found in the gut to outcompete methionine incorporation.

Gut bacteria incubated with 2 mM HPG in 50% fecal supernatant for 2 hours results in saturated HPG incorporation without altering growth **(Fig 1a,b)** or community composition as per 16S rRNA gene sequencing **(Fig 1c)**. The concentration of supernatant has a significant effect on the BONCAT signal (p=0.032, 2-way ANOVA, Tukey’s multiple comparisons test), but HPG concentration does not, once above 1 mM **(Fig 1a)**. LC-MSMS concentrations of methionine in 5 fecal samples from unrelated healthy volunteers.

**Figure 1:**
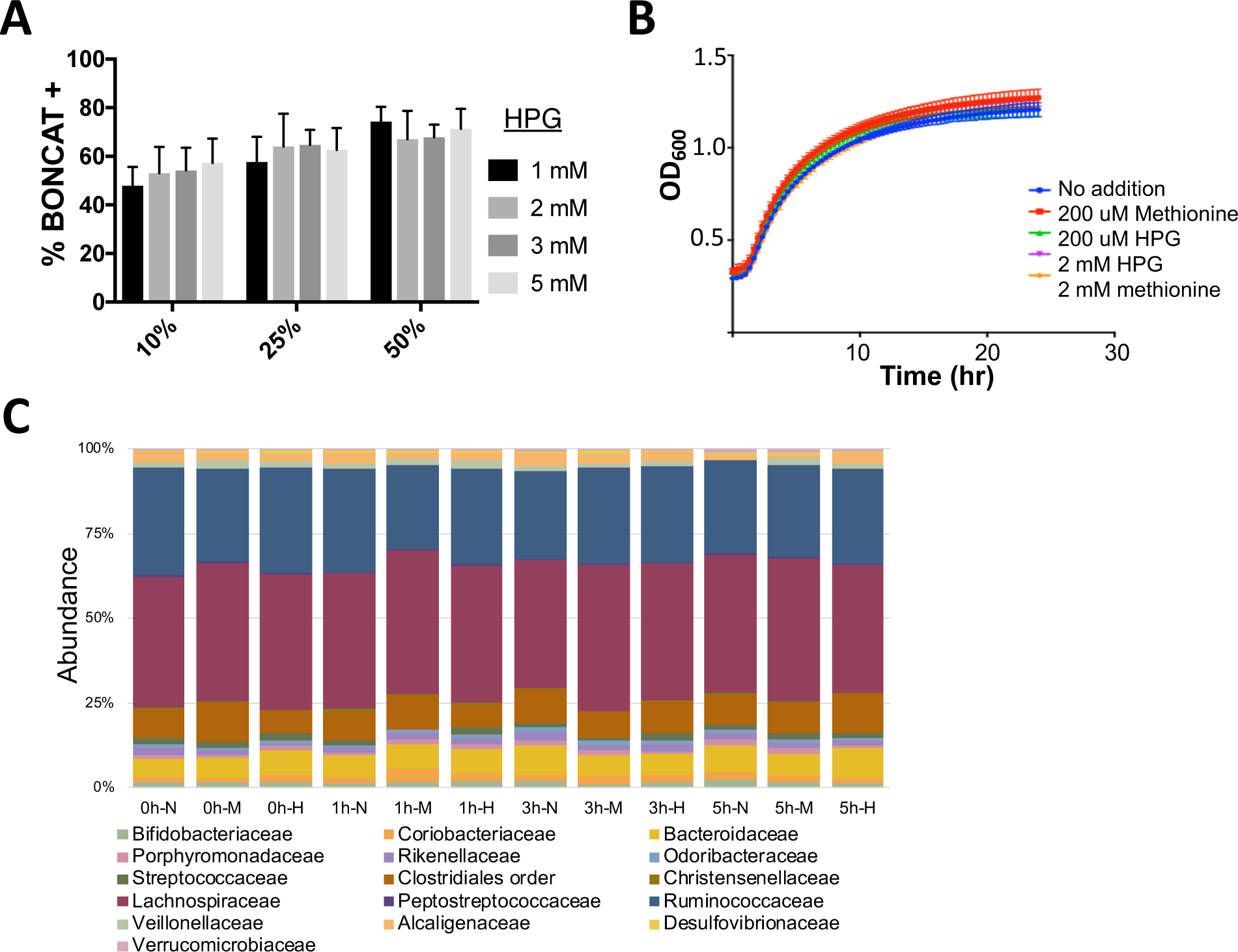
Optimizing in-vitro BONCAT incubation conditions for the human gutmicrobiota. **A)** HPG concentrations between 1 and 5mM tested with varying concentrations of fecal supernatant between 10 and 50% show no significant differences in incubation based on HPG concentration. **B)** Growth curves with HPG, methionine, or neither. Bacteria were diluted 1/10 in 50% supernatant and incubated at anaerobically at 37°C in the dark, with shaking. **C)** Family level 16S rRNA gene sequencing of the gut microbiota with and without HPG or methionine in 50% supernatant for 1, 3, or 5 hours (n=1). Labels indicate time (hours), followed by No addition (N), Methionine (M), or HPG (H).

### Optimizing fluorescence-activated cell sorting (FACS) of BONCAT-labelled bacteria

To link bacterial identity to bacterial activity, we optimized a method to sort the translationally active bacteria (BONCAT+) and sequence with 16S rRNA gene amplification (BONCAT-FACS-Seq). We first verified the click protocol was able to capture all HPG-incorporating bacteria by using *Escherichia coli* in exponential phase as a positive control. Control incubations not containing the fluorophore (Alexa-azide 647) or HPG **(Fig S1ab)** were used to determine the appropriate gating for flow cytometric analysis of the BONCAT+ population, which was 96% BONCAT+ for *E. coli* in exponential phase with glucose supplementation **(Fig S1c)**. For the gut microbiota, similar gating controls are employed in a consistent manner **(Fig S1d)** to determine the BONCAT+ population **(Fig S1e)**. As shown elsewhere, dead bacteria did not uptake HPG[29, 30] **(Fig S1f)**.

We determined that sorting 180,000 events was sufficient to represent the unsorted population. Based on a probability mass function for the binomial distribution, we calculated the theoretical value that if we sort 25,000 events, we would capture 100 bacteria that are present in the initial population at a prevalence of 0.5%. We then sorted a range of events from 50,000 to 1,000,000, confirming we sorted viable bacteria that resulted in amplifiable DNA **(Fig S2ab)**. Sorted samples have a slightly lower alpha diversity than the unsorted samples (**Fig S2c)**, yet still cluster near the unsorted samples based on Bray-Curtis dissimilarity in a principal coordinate analysis with no differences in community composition between sorted and unsorted (R^2^ = 0.027, p = 0.887, PERMANOVA)**(Fig S2d)**. The sorting purity for BONCAT, as determined by re-acquiring the sorted fractions by flow cytometry, however, is lower than sorting with other physiological dyes such as SYBR Green I, with a mean purity of the BONCAT+ fraction at 80% ± 10% **(Table S2)**. The BONCAT- fraction; however, has a higher purity level after sorting at 94.3% ± 9%.

The cell sorting process introduced contaminant DNA into the sorted samples. A negative control was sorted each sorting day, consisting of the cell sorter’s sheath fluid, and was extracted and sequenced alongside the samples. The sheath fluid negative controls contained between 341 – 2,333 reads, mostly assigned to *Pseudomonas*. Thus, to remove sheath fluid contaminants from the samples, we removed reads that were present in the sheath fluid, but absent from the initial, unsorted sample from all samples. This had only a minor effect on the diversity and read count of most samples, the most pronounced effect being in the BONCAT- fraction **(Fig S3)**. Thus, we conclude that BONCAT-FACS-Seq yields generally representative 16S communities of the BONCAT+ and BONCAT- populations and is an appropriate method moving forward.

### Diversity of physiologically distinct fractions of the gut microbiota

To determine the active and damaged members of the gut microbiota, we sorted bacteria based on the BONCAT signal, relative nucleic acid content (HNA and LNA), and membrane damage (PI+). Fresh fecal samples were obtained from ten healthy, unrelated individuals who had not received antibiotics in the past three months, and immediately placed in the anaerobic chamber. Samples were processed and stained anaerobically, using reduced media.

The proportion of each physiological fraction from total quantified cells was determined through flow cytometry. The HNA bacteria average 51.73 ± 17.59 %, BONCAT+ at 49.01 ± 18.54 %, and PI+ at 15.73 ± 14.58 % (**Fig 2a)**. There is a high correlation between the proportion of HNA and the proportion of PI+ bacteria (r = 0.74, p = 0.0136) and a borderline significant negative correlation between the proportion of HNA and the proportion of BONCAT+ (r = -0.62, p = 0.0548) (**Fig 2b)**. 16S rRNA gene amplification and sequencing identified that these physiological cell fractions are distinct from one another and from the whole community. Principal coordinate analysis on pairwise weighted UniFrac distances show that samples cluster strongly by individual stool donor (R^2^ = 0.50712, p= 0.001, PERMANOVA) **(Fig 2c)**, and secondarily based on physiology (R^2^ = 0.15171, p= 0.005, PERMANOVA) **(Fig 2d)**. When broken down by individual, HNA and LNA appear most distinct from one another on the first principal component, and BONCAT- most distinct from the other samples on the second principal component **(Fig 2ef)**.

**Figure 2:**
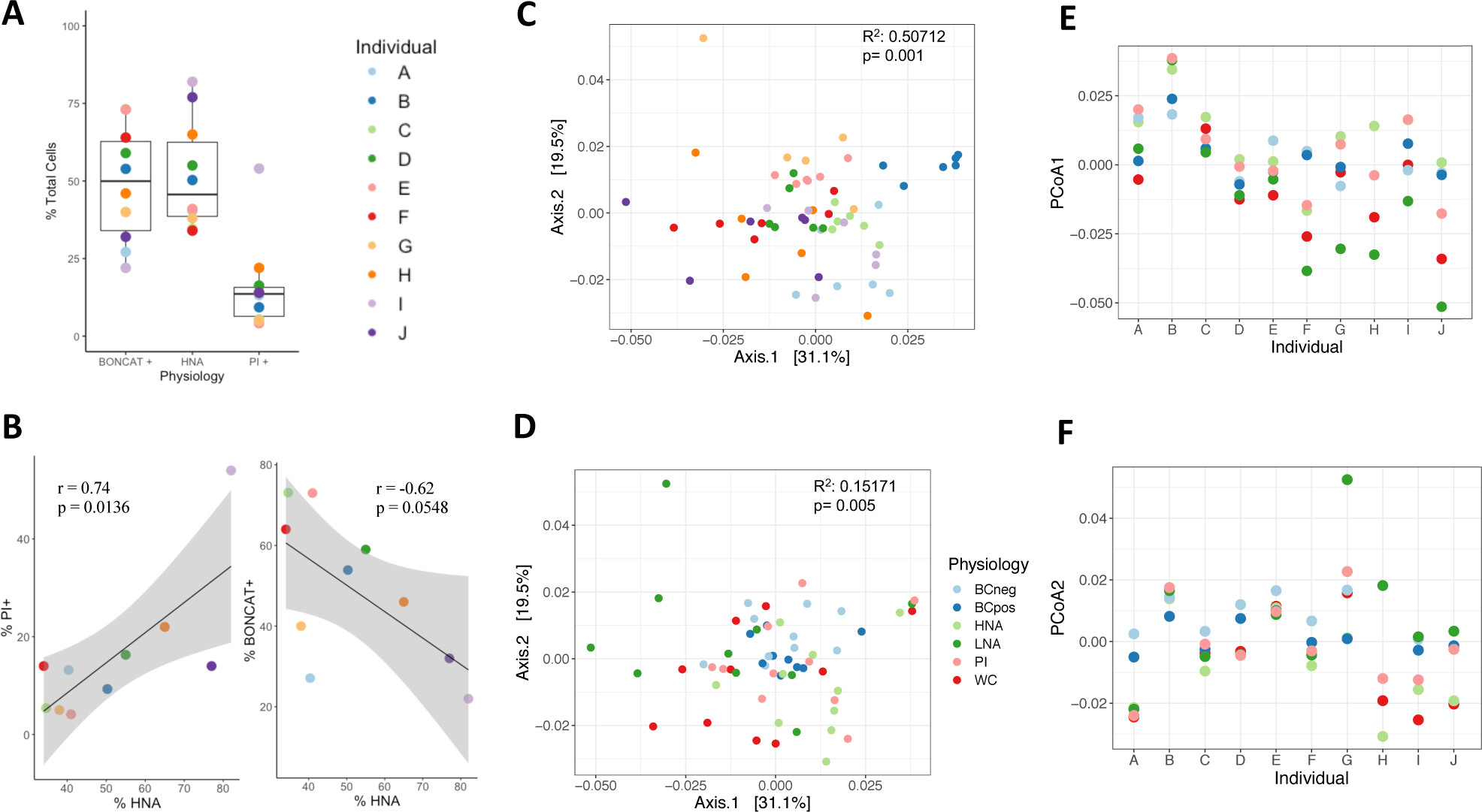
Weighted Unifrac PCoA plots of regularized log transformed data. **(a)** Relative abundance of cells in each physiological fraction (n = 10). **B)** Correlations between the proportion of PI+ bacteria and BONCAT+ bacteria to the proportion of HNA. There is significant clustering by **(c)** individual, and **(d)** physiology. As there is large variation across individuals, the first **(e)** and second **(f)** principle components were plotted by individual to show trends in clustering by physiology. Samples are colour coded by individuals in all panels **A-C** and by physiology in panels **D-F**.

We next set out to determine specific differences between physiological fractions. The HNA and LNA communities are significantly different from one another at the phyla and genus level (p<0.05, PERMANOVA with FDR adjustment) and the HNA is borderline significantly different from the whole community at the phyla level (p = 0.055, PERMANOVA with FDR adjustment) (**Fig 3ab)**. No other fractions were significantly different from one another after FDR adjustment. Phyla and genus level relative abundances are broken down by individual as well **(Fig 3cd)**. To determine specifically which taxa are differentially abundant, we fitted a count regression for correlated observations with the beta-binomial model (Corncob) to every taxa and compared each physiological fraction to the whole community or to their low/high activity counterpart. The HNA fraction is dominated by the Firmicutes, averaging 80.98% of the HNA community compared to 61.28% of the whole community (p = 0.004, corncob). The HNA fraction also contains fewer Actinobacteria compared to the whole community (p = 0.0008, corncob). Similarly, the PI+ fraction resembles HNA more closely than it does the whole community, with significantly more Firmicutes at 78.02% (p = 0.04, corncob) and fewer Actinobacteria (p = 0.017, corncob). Conversely, the LNA fraction is more similar to the whole community, with 44.94% Firmicutes and 51.36% Bacteroidetes. As shown previously in the human gut[10], the HNA and LNA fractions contain the same bacteria (unweighted UniFrac p = 0.6), but present at different relative abundances (weighted UniFrac p = 0.028,), suggesting that nucleic acid content reflects bacterial physiology rather than taxonomy.

**Figure 3:**
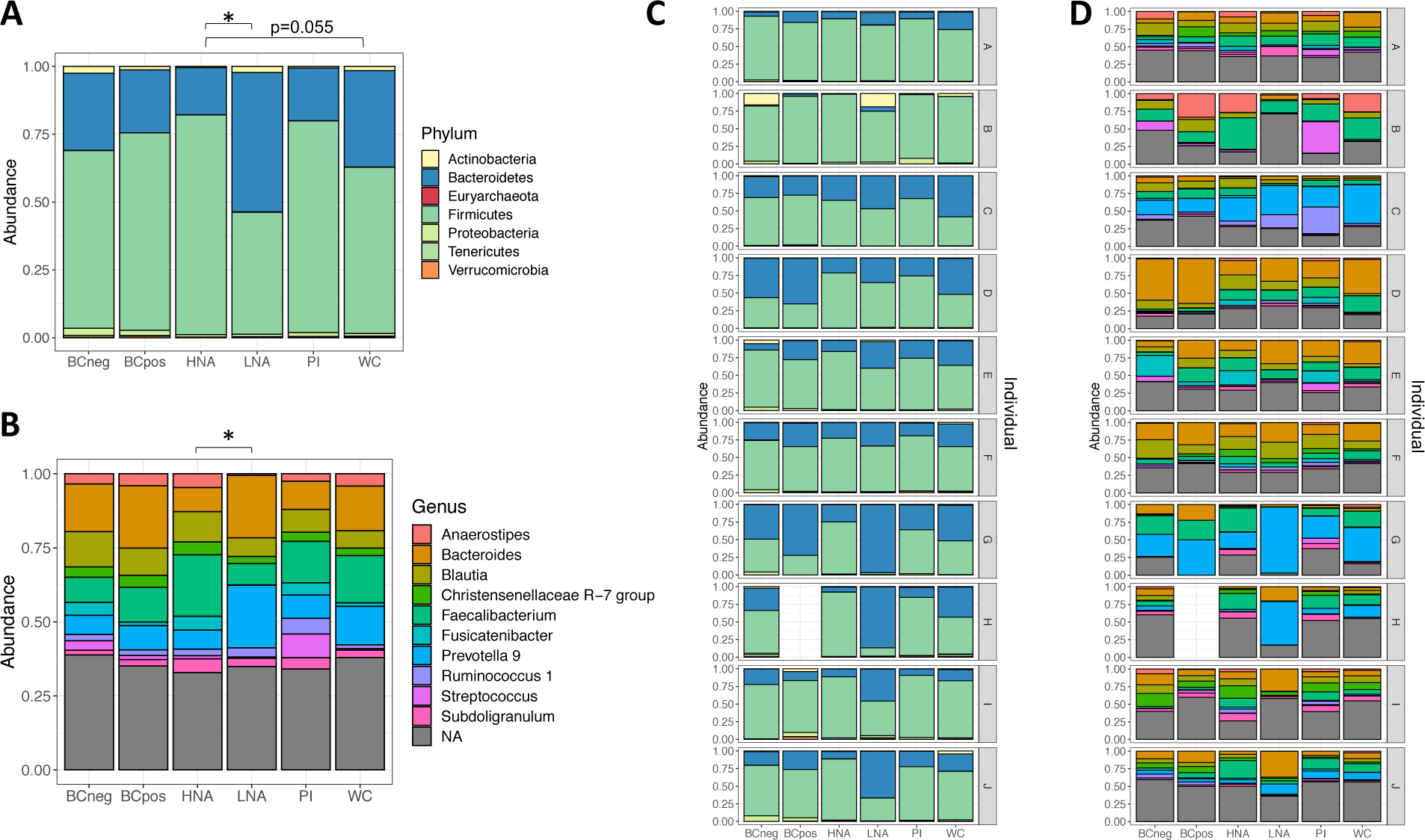
Taxonomic overview of the physiologically distinct subpopulations of the human gut microbiota. Relative abundance at the **(a)** phylum and **(b)** genus level across physiological groups and by individual **(c)** and **(d)**. Pairwise Adonis tests, with FDR adjustment.

Surprisingly, the BONCAT+ and BONCAT- fractions are not significantly different from one another, or from the initial community (PERMANOVA). Still, Proteobacteria are significantly increased in BONCAT- compared to the whole community. While the BONCAT- fraction contains more Proteobacteria in general (p = 0.0069, corncob), the BONCAT+ fraction contain higher abundances of specific lineages within Proteobacteria such as *E. coli/Shigella* (p = 0.045, corncob), as well as differentially abundant *Coprococcus* 1 (p = 3.98e-05, corncob).

### The HNA bacteria of the gut microbiota contain more core taxa

HNA bacteria have previously been characterized as the more active bacteria in a community. Yet we found that the HNA bacteria are not necessarily the protein-producing bacteria, as HNA and BONCAT+ cell fractions are taxonomically different from one another (R^2^ =0.11721, p=0.014, q=0.07 FDR). We hypothesized that HNA and BONCAT+ bacteria each represent different aspects of bacterial activity, but as active members of the community, both should contain more members of the core microbiome (*i*.*e*. taxa commonly present in most people).

Changes in the core bacteria or functional groups have been linked to changes in host phenotype[48–51], suggesting the core microbiome is actively responsible for host phenotype, potentially through the metabolites they produce. Conversely, the less active fraction (BONCAT- and LNA) would contain the transient, environmental bacteria that are unique to the individual and perhaps less adapted to survive in the gut environment[39–42]. Consistent with our hypothesis, the HNA fraction is more similar across individuals than the whole community (p<0.01), or the LNA community (p <0.0001). The LNA fraction is more different across individuals than the whole community (p<0.05, Kruskal-Wallis test with Dunn’s test for multiple comparisons of weighted UniFrac distances) **(Fig 4a)**. However, we found no equivalent differences in the BONCAT communities across individuals.

**Figure 4:**
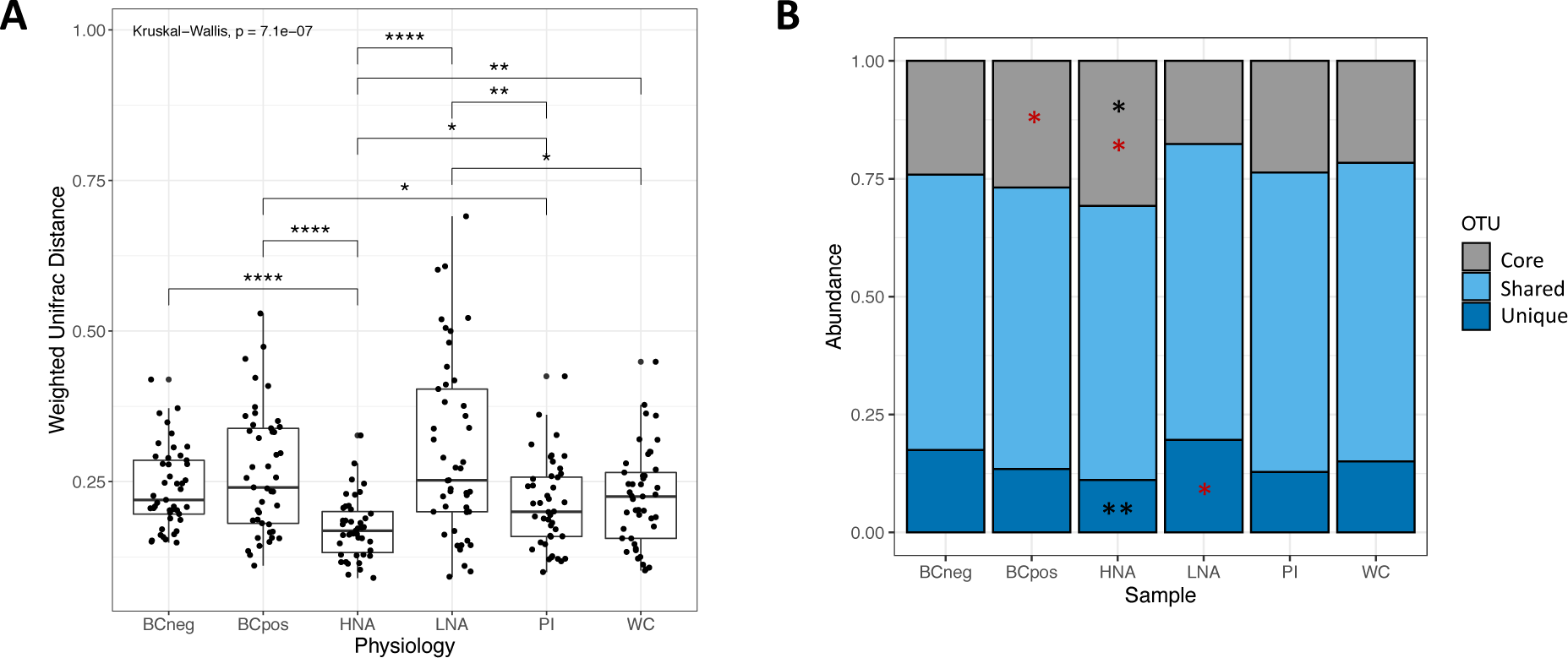
Similarity of physiologically distinct fractions across individuals. **A)** Weighted UniFrac distances for each pair of samples of rlog transformed data. **B)** Distribution of core, unique and shared taxa across physiological fractions. Red stars represent significantly different dispersion, black stars represent significantly differential abundance as per corncob models. \**p* < 0.05, \*\**p* < 0.01, \*\*\**p* < 0.001, and \*\*\*\**p* < 0.0001.

To expand on the idea that the active fractions are more similar across individuals and thus contain more of the common core bacteria, we compared the distribution of core, unique, and shared taxa across the physiologically distinct fractions **(Fig 4b)**. Core taxa are defined as those found in all sampled individuals, unique taxa are defined as those present in only one individual, and shared represent the remaining taxa. With this definition, out of 838 bacterial amplicon sequencing variants (ASVs), 12 (1.4%) were found in all individuals and 477 (57%) bacteria found in only one individual. The remaining 349 bacteria (41.5%) of bacteria are shared across some proportion of individuals. The core taxa are differentially dispersed in HNA and BONCAT+, while also being significantly increased in HNA (p<0.05). HNA also has fewer unique bacteria (p < 0.01, corncob), without a difference in dispersion. These patterns remain when the definition of core bacteria is loosened to ASVs found in 8/10 individuals, where 32 (3.8%) bacteria are considered as core. The enrichment of core taxa and lack of unique taxa in HNA, but not BONCAT, suggests the HNA may be providing more of the core metabolic activity in the gut.

### Physiological information is more sensitive than whole community profiling

Focusing on the active subset of the gut microbiota allows for a finer resolution of changes in activity to be captured than whole community DNA sequencing alone. To demonstrate this, we performed *in vitro* incubations of fecal samples from two individuals with various xenobiotics previously shown to alter the activity of the whole community (through metatranscriptomics), but not bacterial community composition (through 16S rRNA gene sequencing)[10]. We supplemented the BONCAT incubations with digoxin, nizatidine, or glucose, and compared the proportions of the physiological communities to a control incubation without any additions. PCoA plots of the weighted UniFrac distances, focusing on each of the two individuals separately, demonstrate a clear clustering by bacterial physiology (R^2^ = 0.65 and 0.64 respectively, **Table 1**) **(Fig 5ab)**. Treatment alone has no effect; however, when looking at the effect of treatment nested within each physiological fraction, it has the largest effect size at R^2^ = 0.75 (individual 1) and R^2^ = 0.73 (individual 2) **(Table 1)**. To determine specific pairwise effects of xenobiotic treatments relative to the controls in each physiological group in each individual, we compared Bray-Curtis dissimilarities. Specific changes in response to the xenobiotics in each physiological group were inconsistent, and limited by the low number of replicates per sample (n=3). However, a few physiological groups trended towards significance compared to the control incubation. Specifically, in individual 1, the glucose incubation resulted in a change in beta diversity compared to control in the whole community and in the LNA fraction, and digoxin had effects in the PI+ fraction (p=0.1, PERMANOVA). The taxonomic composition of these sorted fractions are in **Figure S4**.

**Table 1:**
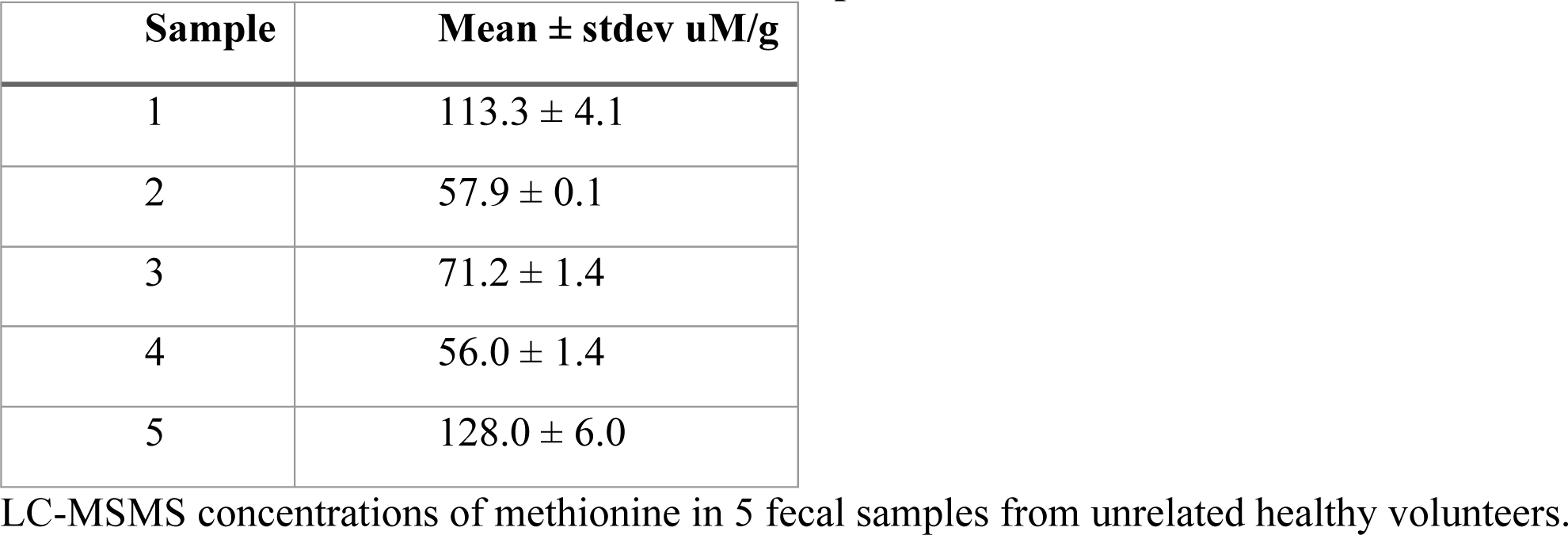
Methionine concentration in stool samples.

| Sample | Mean ± stdev uM/g |
| --- | --- |
| 1 | 113.3 ± 4.1 |
| 2 | 57.9 ± 0.1 |
| 3 | 71.2 ± 1.4 |
| 4 | 56.0 ± 1.4 |
| 5 | 128.0 ± 6.0 |

LC-MSMS concentrations of methionine in 5 fecal samples from unrelated healthy volunteers.

**Table 2:**
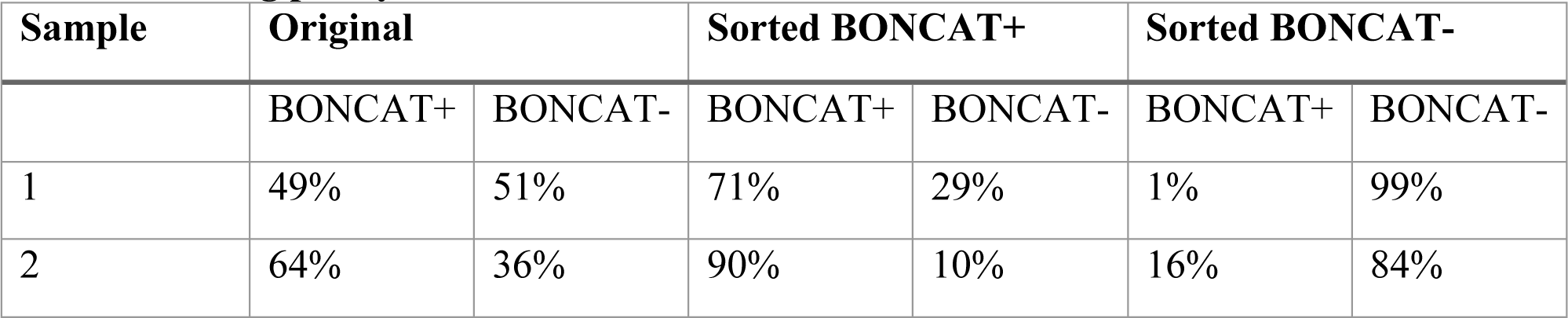
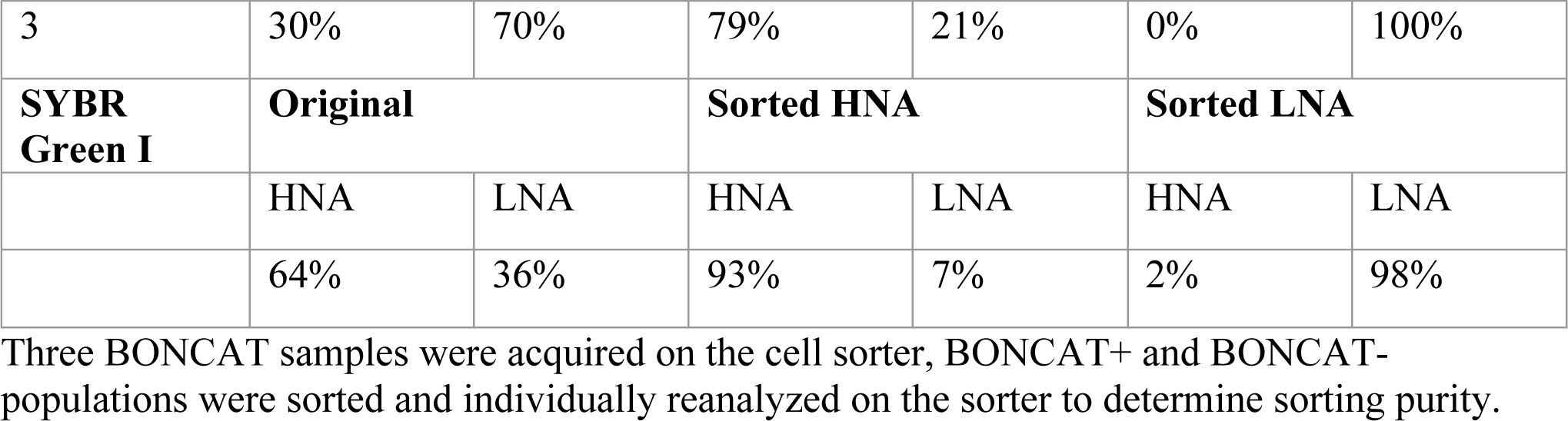
Sorting purity.

**Table 3:** Effect size and dispersion effect of xenobiotics on the gut microbiota.

|  | Individual 1 |  |  |  | Individual 2 |  |  |  |
| --- | --- | --- | --- | --- | --- | --- | --- | --- |
|  | PERMANOVA |  | Dispersion |  | PERMANOVA |  | Dispersion |  |
|  | R<sup>2</sup> | p | F | p | R<sup>2</sup> | p | F | p |
| Physiology | 0.65 | 0.001 | 6.25 | 0.001 | 0.64 | 0.001 | 1.3 | 0.275 |
| Treatment | 0.010 | 0.8 | 0.33 | 0.81 | 0.018 | 0.53 | 0.6635 | 0.582 |
| Treatment %in% Physiology | 0.75 | 0.001 | 0.6574 | 0.861 | 0.73 | 0.001 | 0.69 | 0.825 |

**Figure 5:**
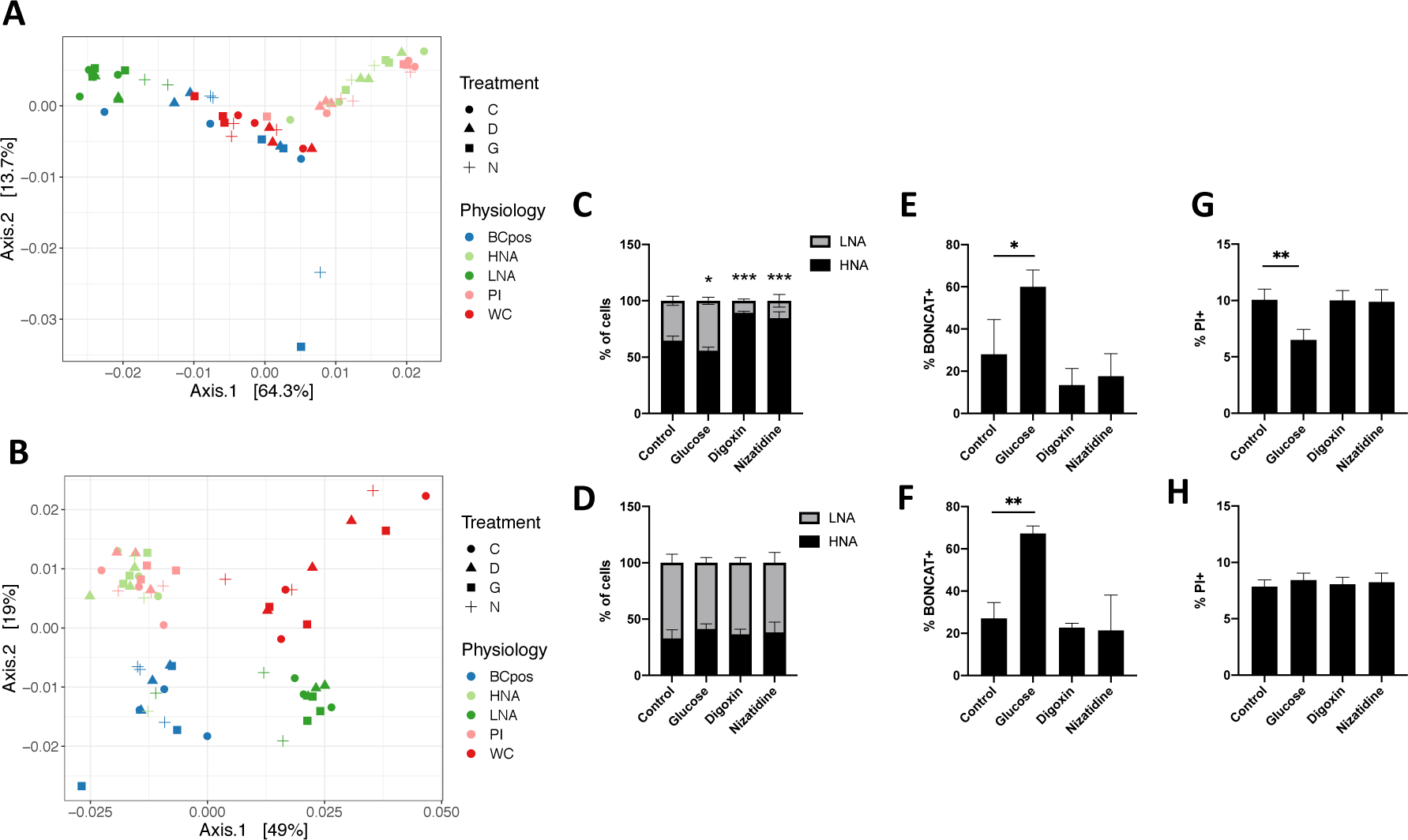
Differences of physiological groups after xenobiotic incubations. PCoA of rlog transformed weighted UniFrac distances broken down by physiology and treatment for **A)** individual 1 and **B)** individual 2. The circle for treatment C represent the control, the triangle for treatment D represents digoxin, the square for treatment G represents glucose, and the plus sign for treatment N represents nizatidine. Cellular relative abundance of **(C)** HNA and LNA, **(E)** BONCAT+, and **(G)** PI+ bacteria in Individual 1. The relative abundance of **D)** HNA and LNA, **F)** BONCAT+, and **H)** PI+ bacteria in Individual 2. Ex-vivo incubations were performed anaerobically at 37°C for 2 hours. N=3 incubation replicates, error bars represent S.D, One-wayANOVA compared to the control group, with FDR correction.

BONCAT-FACS-Seq provides actual cellular abundance data of each of the physiological fractions. The proportion of HNA bacteria decreased in individual 1 in response to glucose (65 ± 4% to 56 ± 3.2%), but increased in response to digoxin (to 89 ± 1.5%) and nizatidine (85 ± 5.5%) **(Fig 5c)**; however, there were no changes in the proportion of HNA and LNA in individual 2 **(Fig 5d)**. Increases in the proportion of HNA bacteria in individual 1 did not correspond to changes in composition, which remained similar to the PI+ fraction (**Figure S4)**. The proportion of BONCAT+ bacteria increased in response to glucose in both individuals (28 ± 17% to 60 ± 7.9% and 27 ± 7.5% to 67 ± 3.5%, respectively) **(Fig 5ef)**. And lastly, the proportion of PI+ bacteria decreased in response to glucose in individual 1 (10 ± 0.96% to 6.5 ± 0.94%) but not individual 2 **(Fig 5gh)**. While the response in HNA/LNA and PI proportions is individual specific, there is a consistent increase in the proportion of BONCAT+ bacteria in response to glucose. Overall, we demonstrate how BONCAT-FACS with or without sequencing may be a faster and less expensive alternative to detect broad-level changes in bacterial activity than metatranscriptomics or whole community profiling.

## Discussion

In this study, we have optimized the BONCAT-FACS-Seq protocol for the human gut microbiota to identify the translationally active bacteria. We then compared the cellular proportions and bacterial diversity of the BONCAT+ and BONCAT- fractions to the high and low nucleic acid containing bacteria (HNA and LNA), as well as the damaged fraction through PI staining. We found that the HNA and PI+ bacteria correlate well both in terms of abundance and diversity. The BONCAT+ and HNA fractions are taxonomically distinct, and as such we suggest they represent two different aspects of bacterial metabolic activity.

The high and low nucleic acid communities have been previously studied in the human gut microbiota. Previous work reported similar proportions of HNA, LNA, and PI in healthy individuals as our study, along with the dichotomous taxonomic distribution where the HNA are dominated by Firmicutes, and Bacteroidetes most abundant in the LNA fraction[10]. This was found both with a DNA-based and an RNA-based dye, and both studies align with ours showing that the HNA fraction is taxonomically distinct from the whole community[10, 18]. Firmicutes have on average smaller genomes than the Bacteroidetes, suggesting genome size is not a factor in this bimodal distribution[52, 53]. In the same study, the Firmicutes were shown to be transcriptionally more active than the Bacteroidetes, and were the first to be damaged upon exposure to a perturbation[10], a characteristic of HNA that has been shown in other studies[54, 55]. The correlations seen with the relative abundance and diversity between HNA and PI suggest that a portion of the HNA bacteria are damaged, or contribute to the pool of damaged bacteria. However, the specific biological mechanisms at play that differentiate the HNA/LNA modality in the human gut remain undefined.

Relative nucleic acid content is thought to be related to the metabolic activity of the cell, as for example, it is supposed that a bacterial cell would contain more RNA when active than when inactive. As most RNA in the bacterial cell is involved in translation, we optimized a complementary technique, BONCAT to specifically probe the production of nascent proteins. BONCAT-FACS is a promising technique that allows for the sensitive, rapid identification of translationally active bacteria from just a short *in vitro* incubation. By adding excess HPG and incubating the bacteria in their own fecal slurry anaerobically, we maintained to *in situ* conditions as much as possible. Thus, we were able to use a defined marker of activity: translation, to compare to an undefined marker of activity: relative nucleic acid content, to determine how much protein production contributes to the activity supposed by relative nucleic acid content.

Both relative nucleic acid content and BONCAT identified approximately half of the gut microbiota as active. This is similar to what has been seen in other studies using stable isotope probing with heavy water (D_2_O), where they found a range in the proportion of active bacteria from 30 to 76%[16]. While the overall average proportion of HNA and BONCAT+ bacteria were similar in this study, their proportions did not correlate within an individual, and these two fractions are taxonomically distinct from one another. This suggests that the HNA bacteria are not necessarily the protein-producing bacteria within the system, and that HNA and BONCAT are two independent pools of bacteria. It is commonly assumed that transcription and translation are tightly coupled[56], so that both RNA and protein activities would be correlated and a relationship between BONCAT+ and HNA cells would exist. However, it has recently been shown that this tight coupling is not always the case, and specifically, there exists a substantial lag between transcription and translation across most Firmicutes[57]. This is similar to what we are seeing in the human gut microbiome: there is no clear relationship between BONCAT+ and HNA cells, consistent with the notion that there is large variation in the relationship between transcription and translation across bacteria.

The HNA fraction is the physiological fraction most similar across individuals, containing more core taxa and less unique taxa than the whole community. Thus, the HNA cells may provide the bulk of the common cellular functions performed by the gut microbiota, containing the conserved functionality required for members of the gut microbiome to exist in the intestinal milieu. In this sense, the HNA fraction is metabolically active and still functionally relevant, but less resistant to damage than the BONCAT+ fraction.

The translationally active bacteria (BONCAT+) are not taxonomically distinct from the whole community or from the non-translating fraction (BONCAT-). Upon glucose addition, the proportion of BONCAT+ bacteria increased significantly, more than doubling in relative abundance. This suggests that most of the bacteria present in the gut have the potential to become translationally active, and that BONCAT with or without supplementations is able to differentiate between the actual and potential activity of the gut microbiota. Previous studies have suggested that approximately 20% of the gut microbiota is dormant, with 15% of the gut microbiome containing sporulation homologues[43]. While the non-translating fraction determined in this study is larger than what has been predicted to be dormant, it may represent a bacterial reservoir; a spectrum ranging from dead, damaged, fully dormant, to a slower rate of protein production not captured with the incubation time used here. The lack of distinction between the BONCAT + and BONCAT- fractions could possibly be due to the leakiness in the sorting, but with purity levels ranging around 80%, only minor changes in the diversity between BONCAT+ and BONCAT- would be missed. As 16S rRNA gene sequencing rather than shotgun metagenomics was performed on the sorted fractions, differences might exist between the BONCAT + and BONCAT- fractions and the whole community at a higher taxonomic resolution, afforded by methods such as shotgun metagenomics.

Adding physiological information to sequencing data provides more sensitivity to subtle changes in the gut microbiota than whole community diversity changes alone. The changes in the proportions of these physiological fractions in response to various drugs or glucose demonstrate the utility of a rapid method to determine changes in bacterial activity that occur before changes in taxonomic composition. The substantial increase in the proportion of BONCAT+ bacteria after glucose addition is in line with previous work[15, 16], and demonstrates how the majority of the gut microbiota can be stimulated, differentiating between the realized potential of the community rather than the theoretical upper limit. BONCAT is a sensitive method to detect changes in the active fraction of the human gut microbiota, but the lack of changes in diversity of these fractions highlights the heterogeneity of activity in microbial communities seen elsewhere[36]. Changes in the proportion of HNA bacteria in response to certain xenobiotics were not as consistent as changes in the BONCAT+ fraction, but could represent an increase in stress response based on the correlation with damaged bacteria.

We believe BONCAT is a suitable method to study the translationally active members of the human gut microbiota. The limitations of BONCAT are well described in this comprehensive review[19]. Specific to this study, the combination of BONCAT with FACS-Seq remains an area to further optimize, as our sorting efficiency was lower than sorting SYBR-stained cells. It is possible that bacterial aggregates with a mixture of positive and negative cells are being sorted together as BONCAT+. Further optimization of the BONCAT-FACS protocol, for example to allow for double positive sorting could further enlighten the HNA/LNA distribution and its relationship to translational activity. Other measurements of activity, such as replication or transcription, are amenable to the gut microbiota incubation and click protocol, and would help further characterize the contribution of relative nucleic acid content as a physiologically marker of activity.

### Conclusion

In this study, we compared two broad indicators of bacterial metabolism using cell sorting methods: relative nucleic acid content and translational activity. Both markers identify approximately half of the community as active, yet are distinct from one another. Thus, the HNA bacteria are not necessarily the protein-producing bacteria, and these two fractions represent distinct types of metabolism and activity. By focusing on the active subcommunities of the gut microbiota, we can more sensitively detect changes to perturbations than looking at the whole community alone. We hope this work lays the groundwork for using bulk-activity measurements to study how bacteria are able to change their physiology in response to various perturbations.

## Declarations

### Ethics approval and consent to participate

Informed consent was obtained from all volunteers. The study was approved by the McGill Ethics Research Board (REB #A04-M27-15B), Montreal, QC, Canada.

### Consent for publication

Not applicable.

### Availability of data and materials

Bacterial 16S rRNA gene sequencing data can be accessed on the SRA database, accession number XXXX. Code related to the analysis has been deposited in GitHub (https://github.com/MTaguer).

### Competing interests

The authors declare that they have no competing interests.

### Funding

This work was supported by a Natural Sciences and Engineering Research Council (NSERC) fellowship awarded to MT, the Canada Research Chair program (950-230748 X-242502), a Canadian Institutes of Health Research (CIHR) transition grant to CFM (PJT-149098), and the Azrieli Global Scholar Program from the Canadian Institute for Advanced Research to CFM.

### Authors’ contributions

MT designed, performed, analyzed and interpreted all the experiments performed in this manuscript. CFM obtained funding and helped design experiments and with data interpretation. BJS aided in writing the manuscript. All authors edited the manuscript and approved the final draft.

## Acknowledgements

The authors would like to thank members of the Maurice lab in their aid in editing parts of the manuscript. As well, the authors would like to thank Josep Gasol, Marta Sebastian, and Marga EC for their useful advice on the BONCAT procedure. The flow cytometry work was performed in the Flow Cytometry Core Facility for flow cytometry and single cell analysis of the Life Science Complex and supported by funding from the Canadian Foundation for Innovation.

## Supplemental Figures

**Supplementary Figure 1:**
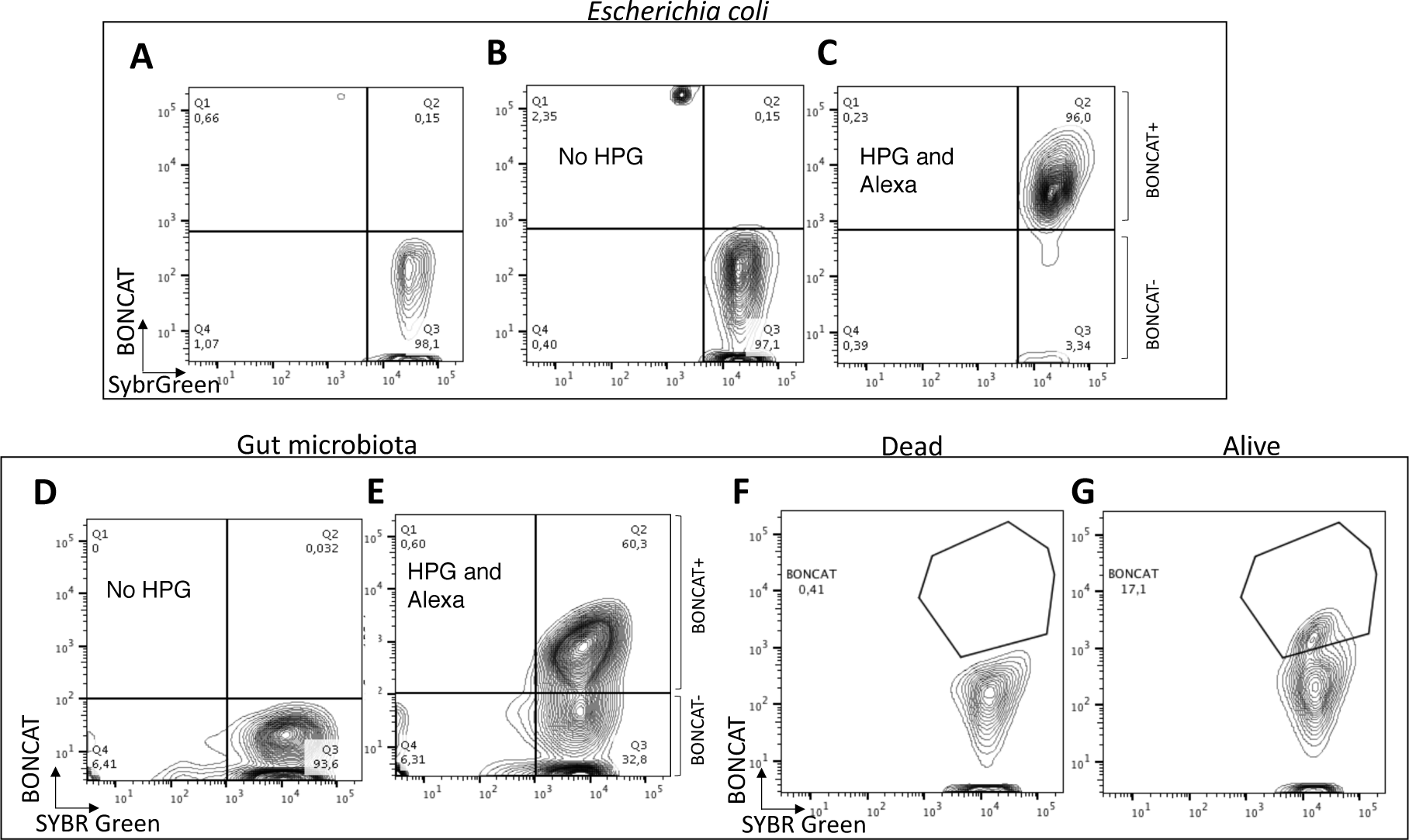
Fluorescently activated cell sorting (FACS) of BONCAT samples. Escherichia coli samples incubated **A)** with HPG but no Alexa and **B)** no HPG but with Alexa as gating controls, for **C)** E. coli incubated with HPG and clicked with Alexa. The gut microbiota **D)** incubated without HPG but with Alexa as a gating control for analysis with both HPG and Alexa in **E)**. Ethanol-fixed microbiota incubated with HPG and Alexa **F)** do not incorporate HPG compared to the same sample not fixed with ethanol **G)**. Q1 is SYBR Green negative and BONCAT positive, Q2 is SYBR Green positive and BONCAT positive. Q3 is SYBR Green positive and BONCAT negative. Q4 is SYBR Green negative and BONCAT negative.

**Supplementary Figure 2:**
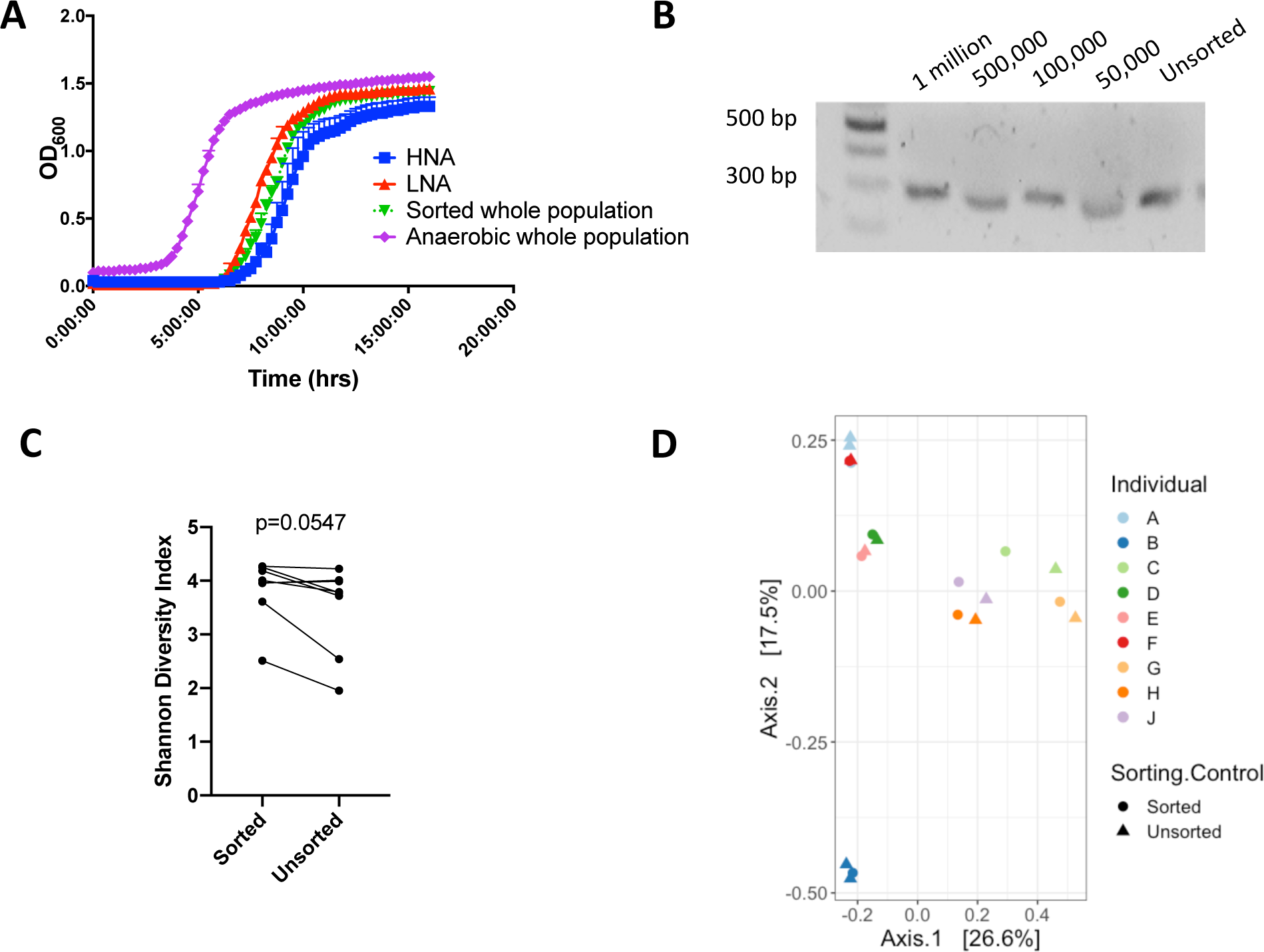
Optimization of sorting to recapitulate the diversity of the initial population. **A)** Anaerobic growth curves of sorted samples compared to an unsorted sample kept in anaerobic conditions. **B)** Between 50,000 and 1,000,000 bacterial events were sorted and 16S rRNA gene amplification by PCR of sorted and unsorted samples. **C)** Shannon’s diversity index of 16S rRNA gene sequencing of sorted and unsorted samples show a borderline significant decrease in alpha diversity (n=8, paired Wilcoxon rank test). **D)** Bray-Curtis PCoA of 16S rRNA gene sequencing results of sorted and unsorted samples for each individual.

**Supplementary Figure 3:**
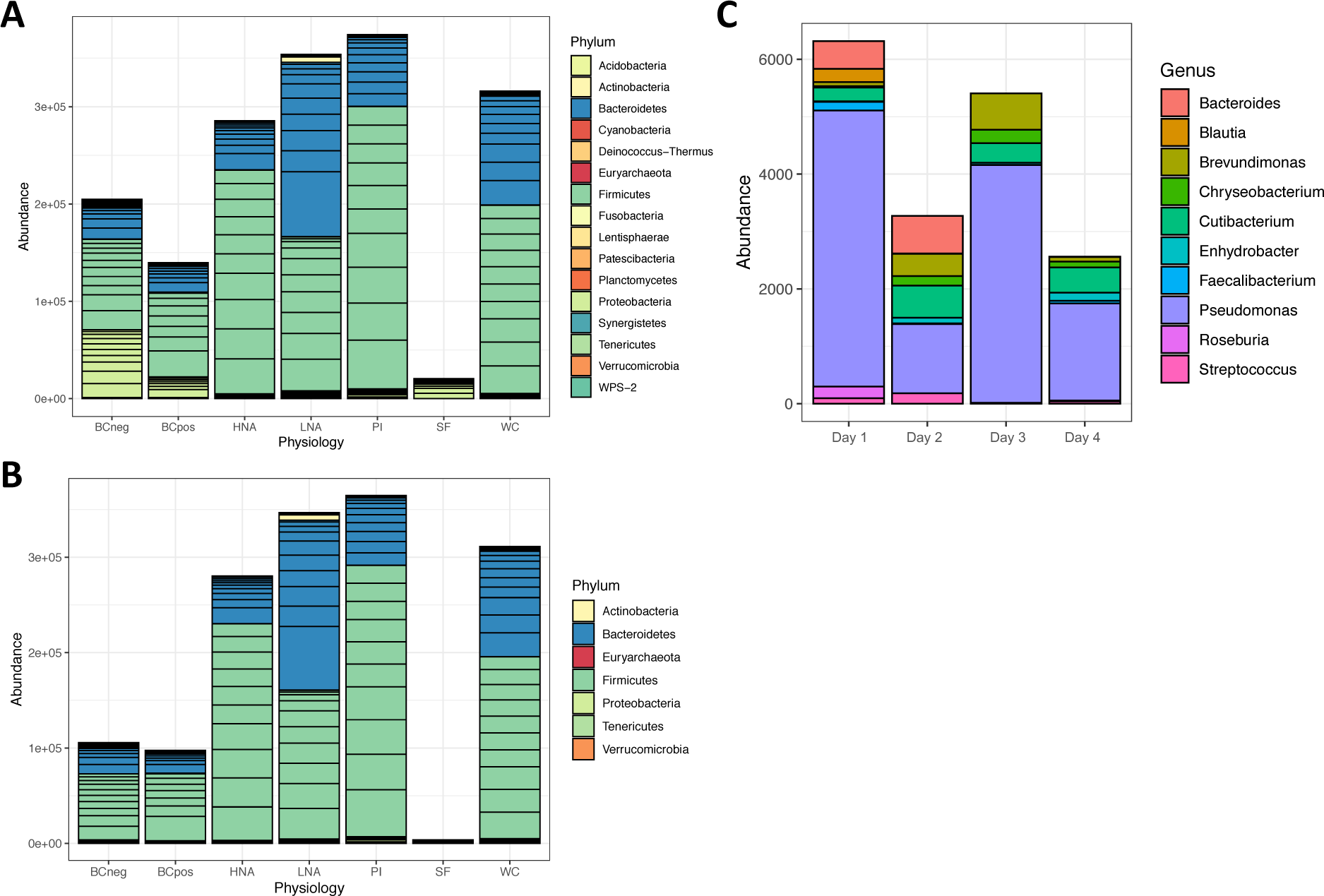
Filtering based on negative control has minimal effect. **A)** Phyla level read counts before any filtering was performed. **B)** Phyla level read counts after taxa present in the sheath fluid but absent from the unsorted sample (DNA) removed from all samples, as well as prevalence-based filtering. **C)** Genus level read count of the sheath fluid samples on the four sorting days. Note the differences in scale of abundance for panel C relative to panels A and B.

**Supplementary Figure 4:**
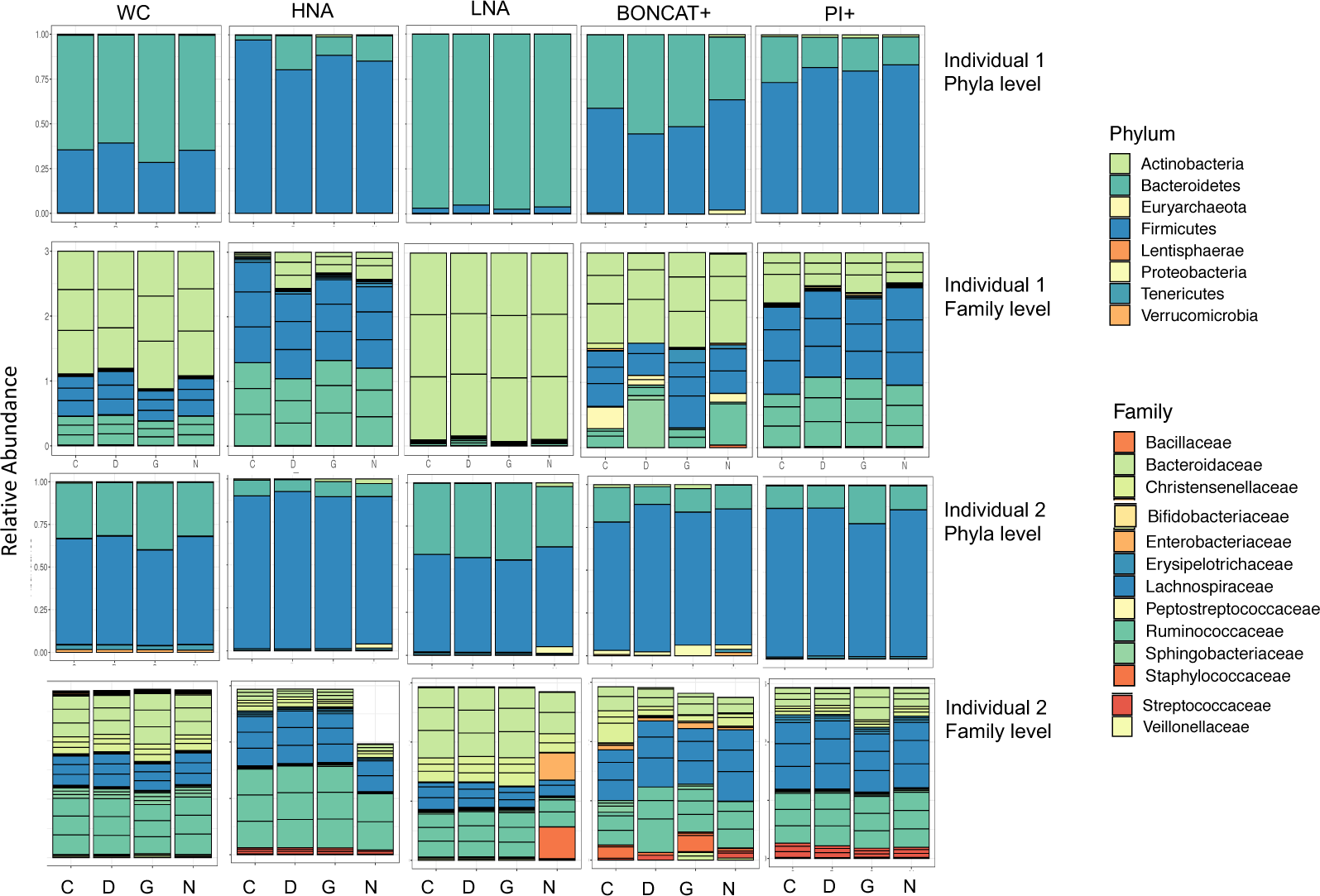
Relative abundances of phyla and family level taxa from 16S sequencing both individuals. Sum averages from each of the n=3 incubation replicates for each xenobiotic.

## Notes

### Competing Interest Statement

The authors have declared no competing interest.

